# Granulysin antimicrobial activity promotes dormancy in *Mycobacterium tuberculosis*

**DOI:** 10.1101/2024.09.27.615427

**Authors:** Sarah Schmidiger, Erin F. McCaffrey, Jan M. Schmidt, Owais Abdul Hameed, Max Mpina, Anneth Tumbo, Elirehema Mfinanga, Frederick Haraka, Hellen Hiza, Mohamed Sasamalo, Jerry Hella, Michael Walch, Jacques Fellay, Sébastien Gagneux, Klaus Reither, José M. Carballido, Ainhoa Arbués, Damien Portevin

## Abstract

Human tuberculosis (TB) caused by *Mycobacterium tuberculosis* (*Mtb*) remains a global public health threat. Granulomas constitute a hallmark of TB pathogenesis that can clear, contain or exacerbate an infection. Containment is exploited by *Mtb* as a hideout to persist in a dormant, antibiotic-tolerant state only to resuscitate upon immunosuppression. The immune determinants of a granulomatous response driving *Mtb* persistence remain elusive. We here combined an *ex vivo* granuloma model with peripheral blood mononuclear cell (PBMC) specimens from TB patients and a high-dimensional mass cytometry (CyTOF) approach to shed light on the immune factors prompting *Mtb* dormancy. Compared to healthy controls, patient-derived *ex vivo* granulomas rapidly force *Mtb* to adopt a dormant-like state; an observation that correlates with the presence of activated innate (-like) cytotoxic lymphocytes. We further demonstrate that *Mtb* dormancy is induced by direct exposure to granulysin, thereby unravelling an immune escape mechanism to cytotoxic lymphocyte activity.

## INTRODUCTION

The immunopathogenesis of tuberculosis (TB), a deadly disease of global prevalence^1^, revolves around the formation of granulomas. Granulomas are organized structures of immune and non-immune cells that form at an infection focus. More than a mere physical containment that ideally clears *Mycobacterium tuberculosis* (*Mtb*) infection, granulomas may represent a niche for *Mtb* persistence, dissemination and ultimately transmission^2, 3, 4^. Far more dynamic than originally postulated, granulomas constitute the ground for a highly complex host-pathogen interplay^5^ that has evolved during millennia. *Mtb* has adopted several strategies to survive host responses and antibiotic treatment^6, 7^. One such strategy is to persist within granulomas by entering into a metabolically dormant state. Dormant mycobacteria are characterized by cytosolic lipid inclusions made of triacylglycerols, loss of acid-fastness, a stagnating metabolism and lack of bacterial division, contributing to an elevated tolerance to antibiotics^8^. Indeed, the presence of lipid-rich bacilli in patient sputum samples has been associated with treatment failure and disease relapse^9, 10^. The molecular mechanisms driving *Mtb* into a dormant state are complex and encompass nutrient and oxygen starvation, as well as oxidative stress^11^. However, whether immune mechanisms beyond environmental or macrophage-specific response play a role in *Mtb* dormancy remains unclear.

Research in TB has studied the role of macrophages extensively; these are the first cell type thought to encounter *Mtb* and to provide a niche for the intracellular replication of the bacteria^12, 13^. The beneficial role of CD4 T cells delivering IFN-γ within ensuing granulomatous responses to *Mtb* infection is well supported by the increased susceptibility of HIV patients^14^ and children displaying Mendelian susceptibility to mycobacterial infections^15, 16^. More recently, cytotoxic lymphocytes have gained attention, with several articles investigating their contribution to immunity against *Mtb* in humans^17, 18^ and non-human primates^19, 20, 21^. These previous studies mainly focused on assessing the immune response’s effects on *Mtb* load, while the impact on the bacilli’s metabolic state has largely been neglected.

We previously demonstrated the relevance of 3D *in vitro* granulomas to reproduce the immune pressure driving dormancy induction and resuscitation of *Mtb* following exposure to TNF-α antagonists, as observed *in clinico*^22, 23^. We here aimed to dissect the host cellular drivers of *Mtb* dormancy within *ex vivo* granulomas using PBMC specimens from a Tanzanian cohort of patients with clinical TB and healthy controls. Using a high dimensional mass cytometry approach, we find patient granulomatous responses to swiftly induce *Mtb* dormancy and identify activated, cytotoxic CD56^+^ lymphocytes as the main drivers. Finally, we demonstrate that sub-mycobactericidal concentrations of granulysin, an antimicrobial peptide, drive *Mtb* into a dormant-like phenotypic state.

## RESULTS

### Patient-derived *ex vivo* granulomas promote a dormant-like *Mtb* phenotype

The fate of TB granulomas is dictated by an intricate interplay of various immune cells and *Mtb*. Combining a granuloma model^24^ with peripheral blood mononuclear cells (PBMCs) of Tanzanian healthy controls and individuals with infectious, clinical TB^25^ (patients), we evaluated how *Mtb* growth and dormancy are affected by immune cells with past exposure and subsequent immune memory to mycobacteria (**Fig. 1a, Supplementary Table 1**). We assessed the differential levels of immune memory to mycobacteria within each PBMC specimen by quantifying the frequency of IFN-γ- producing CD4 T cells responsive to stimulation with an *Mtb*-specific peptide pool or protein extracts (**Fig. 1b, Fig. S2a-b**). Healthy controls showed significantly lower responses to *Mtb*-specific peptides ESAT-6/CFP-10/TB7.7 than patients. In line with previous reports demonstrating that CD4 T cell reactivity to ESAT-6/CFP-10 derived peptide pools is only prominent in about 60% of patients^26, 27^, our patient samples were unevenly reactive to *Mtb*-specific peptide pool stimulation. However, stimulation of samples with *Mtb* soluble-cell wall proteins (SCWP) and whole-cell lysate (WCL) elicited comparable frequencies of IFN-γ^+^ CD4 T cells between the two groups (*p* = 0.15 and *p* = 0.10, respectively). Given the high degree of conservation between the genomes of *Mycobacterium bovis* BCG or environmental mycobacteria with *Mtb*^28^, our data suggest a high level of pre-exposure to mycobacteria also in the healthy controls. Upon *ex vivo* infection of the respective PBMC specimens with *Mtb* H37Rv, the appearance and kinetics of cellular recruitment and aggregation assessed by bright field microscopy did not differ systematically between the two groups (**Fig. S1**). Yet, inter-donor variability was observed as previously reported^29^. In addition, the range of bacterial load retrieved from *ex vivo* granulomas eight days post-infection did not differ statistically between healthy controls and patients (**Fig. 1c**). The prevalence of *Mtb*- or mycobacteria-specific CD4 T cells within the PBMC specimens did not correlate with the level of the *Mtb* load recovered from *ex vivo* granulomas (**Fig. S2c**). Strikingly, bacilli retrieved from patient-derived *ex vivo* granulomas 24 hours post-infection already displayed a marked dormant-like phenotype – reflected by the accumulation of intra-cytosolic lipid inclusions and loss of acid-fastness – in around 45% of bacilli (median 46.3% vs. 33.8% in healthy controls, **Fig. 1d**). Such levels of lipid-rich *Mtb* were only reached eight days post-infection in healthy control-derived granulomas (**Fig. S2d**). The prevalence of pre-existing *Mtb*-specific CD4 T cells did not correlate with the fraction of dormant-like *Mtb* recovered one day post-infection (**Fig. S2e**), suggesting that independent cell subsets or functions promote this rapid metabolic shift in *Mtb*. Taken together, our data suggest that frequent mycobacterial pre-exposure translates into a marked immune memory to mycobacterial antigens in Tanzanian individuals, independently of their TB history. While comparable in their capacity to control *Mtb* growth, patients’ specimens – unlike healthy controls’ – produced a unique immune environment forcing *Mtb* to promptly adopt a dormant-like state.

**Figure 1.**
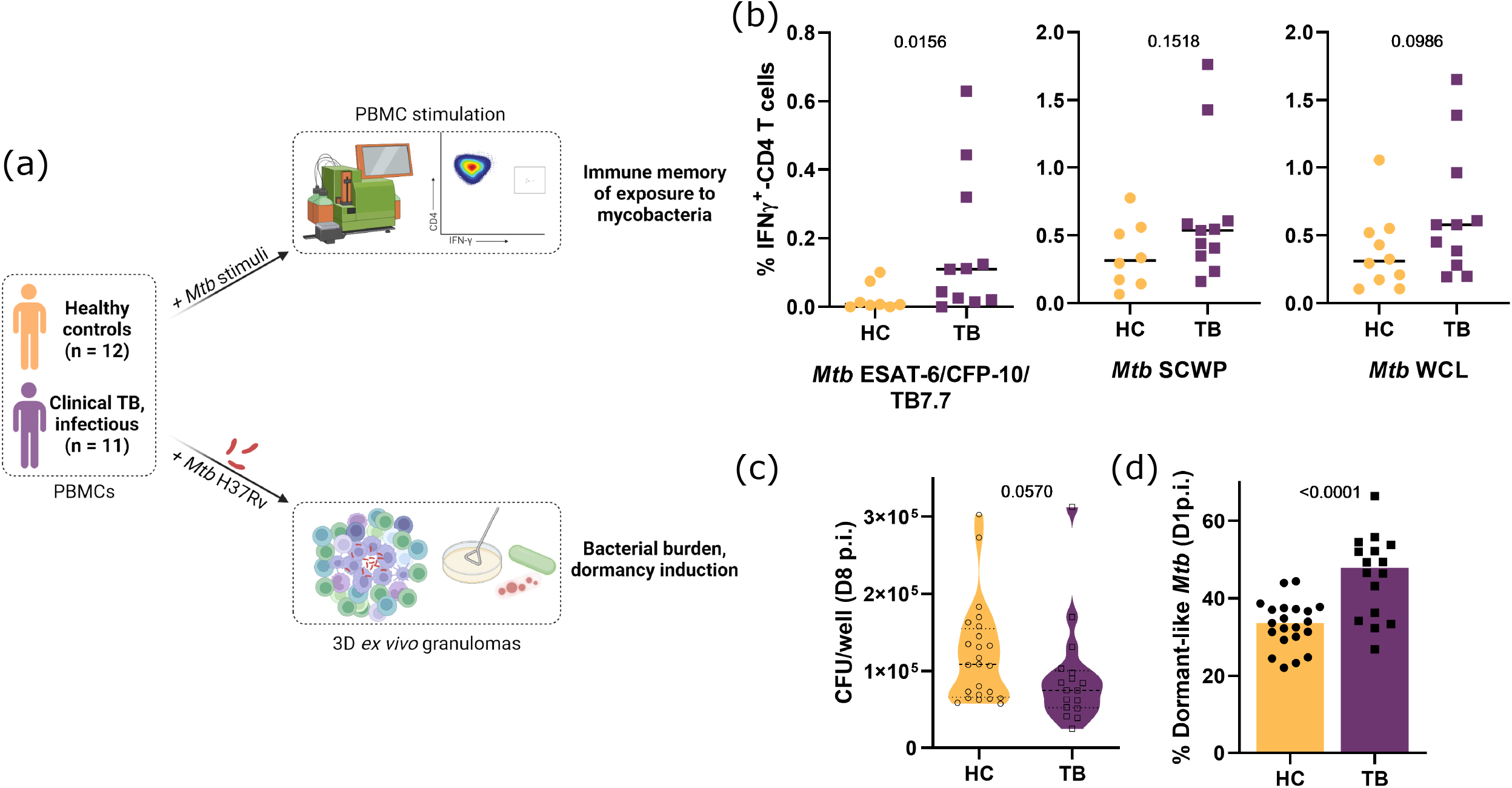
Patient-derived *ex vivo* granulomas promote a dormant-like *Mtb* phenotype. (a) Schematic overview of the study design. Peripheral blood mononuclear cells (PBMC) from healthy controls (HC) or patients diagnosed with infectious clinical TB (TB) were either exposed to *Mycobacterium tuberculosis* (*Mtb*)-derived stimuli prior flow cytometry analysis (b) or infected with *Mtb* H37Rv, prior embedding within an extra-cellular matrix for 3D ex-vivo granuloma formation and assessment of the indicated microbiological read-outs (c-d). (b) Dot plot displaying background subtracted frequency of IFNγ-producing CD4 T cells measured by flow cytometry after overnight stimulation with ESAT-6/CFP- 10/TB7.7 peptide pool, *Mtb* soluble-cell wall proteins (SCWP) or *Mtb* whole-cell lysate (WCL); lines indicate median. (c) Bacterial load in colony forming units (CFU), eight days post-infection (p.i.). (d) Frequency of dormant-like *Mtb* based on auramine-O/Nile red dual staining, one day p.i.; bars depict the median. All *p* values represent Mann-Whitney tests.

### CyTOF-based immune profiling of *ex vivo* granulomas characterizes 25 host cell subsets further subdivided into 640 host cell phenotypes that readily discriminate responses of TB patients from those of healthy controls

To identify the specific host cell subsets promoting the observed metabolic shift in *Mtb*, we employed a high-dimensional mass cytometry by time-of-flight (CyTOF) approach and analyzed the combinatorial expression of 41 markers (**Supplementary Table 2**) in *ex vivo* granulomas 24 hours post-infection. FlowSOM clustering based on 28 canonical markers identified 19 cell subsets of lymphoid and 6 of myeloid origin (**Fig. 2a-b**). As visualized by a uniform manifold approximation and projection (UMAP), *ex vivo* granulomas were mainly composed of T cells (mean frequency: 80.9%) (**Fig. 2c**). NK cells and B cells were similar in frequency (9.8% and 8.5%, respectively) and myeloid cells – which encompassed monocytes, macrophages and dendritic cells – constituted the smallest fraction (0.7%). We next sought to determine whether the frequency of individual cell subsets would correlate with the faculty of the granulomatous response to promote the dormant-like features of *Mtb* bacilli. We detected weak to moderate, positive and inverse correlations of immune cell subset frequencies with the proportion of lipid-rich bacilli; yet, none reached statistical significance (**Fig. S3**). Consequently, we further characterized the phenotypic traits of the individual cell subsets based on the expression of 28 functional markers (**Fig. 2d, Supplementary Table 2**). For each identified cell subset, we determined the frequency of positive cells for each individual functional marker, rendering 700 potential “cell subset-functional marker” combinations (hereafter referred to as “host cell phenotypes”). This approach does not account for collinearity between independent variables; however, principle component analysis (PCA), applied thereafter, is a dimensionality reduction tool that can manage multicollinearity^30^. We used 640 host cell phenotypes that were quantifiable across all samples to perform a PCA. Due to the multitude of highly covariant variables used to calculate the PCA, the variance accounted for by the first two PCs (11.8% and 8.1%) are relatively low. Yet, the first principle component (PC1) could readily separate patient and control samples (**Fig. 2e**), suggesting that CyTOF-captured functional responses were substantially distinct between the two groups. PC2 revealed further variance, especially within the patient group. Taken together, this high dimensional characterization of granulomatous responses to *Mtb* infection revealed substantial profile differences that primarily depend on the individuals’ disease status.

**Figure 2.**
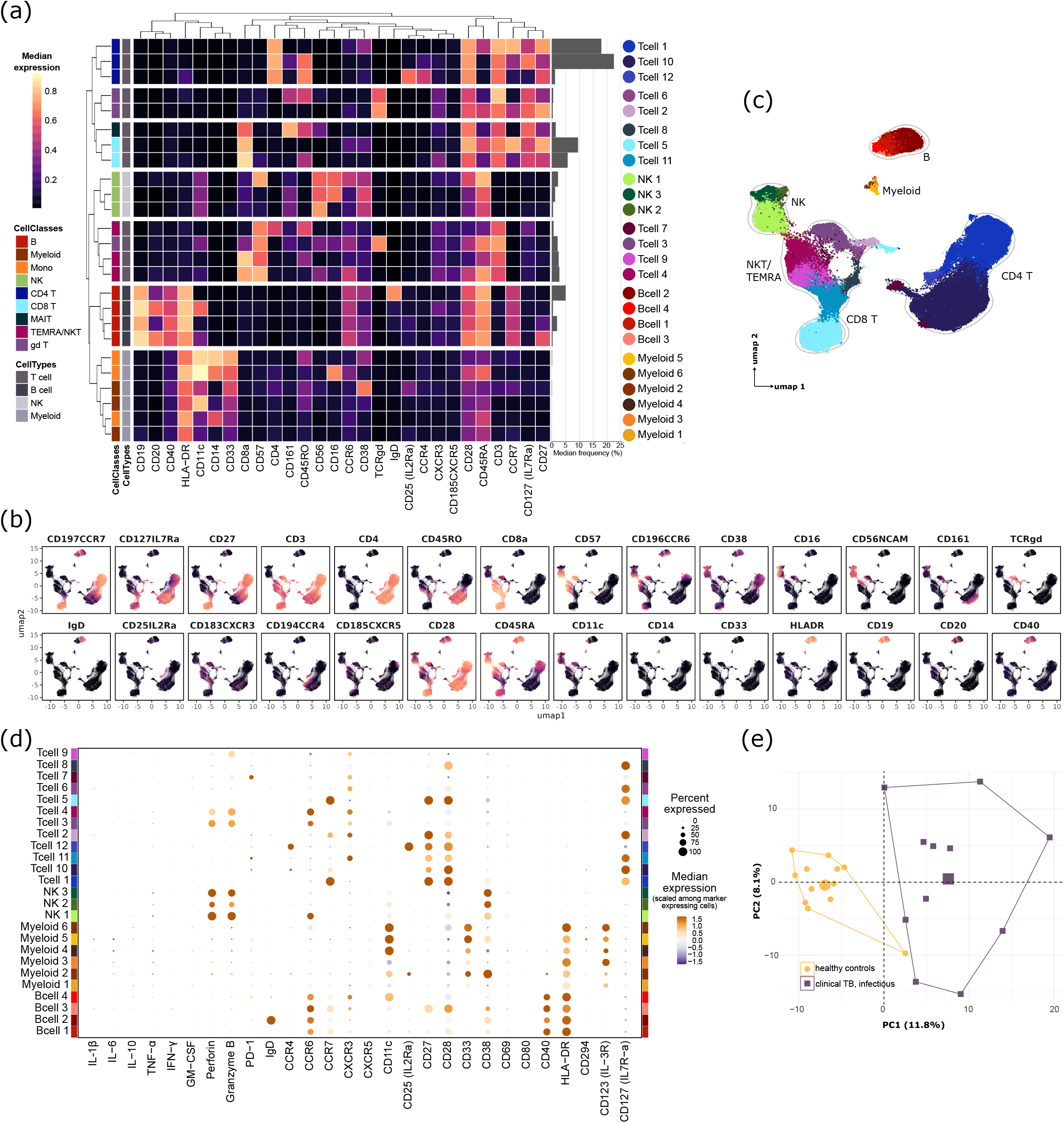
CyTOF-based immune profiling of *ex vivo* granulomas characterizes 25 cell subsets further subdivided into 640 host cell phenotypes that readily discriminate responses of TB patients from those of healthy controls. (a) Heatmap of the median expressions of FlowSOM clustering markers with a horizontal bar plot representing the median frequency of the respective cell subsets specified by adjacent colored circles in the legend. UMAP visualization colored by (b) median expression of clustering markers, and (c) FlowSOM cluster identity. (d) Percent expressed and median expression level of functional markers characterizing the 25 subsets (host cell phenotypes, n=700). (e) Principal component analysis on the host cell phenotype frequencies common to all *ex vivo* granulomas from study participants (n=640), colored by disease group.

### Cytotoxic, activated NK and (unconventional) T cells are significantly enriched in patient-derived granulomas

To define which of the 640 host cell phenotypes significantly drive the immune profiles that distinguish TB patients from healthy controls in the PCA, we evaluated the effect size and statistical significance of differential host cell phenotypes (**Fig. 3a**). At a statistical significance threshold of *p* < 0.05, we detected 14 host cell phenotypes enriched in healthy control responses and 10 in patients’ (**Supplementary Table 3**). Patient-derived profiles were enriched for activated, cytotoxic T and NK cell phenotypes, as well as for CD38-expressing myeloid cells. Applying a stringent threshold of *p* < 0.01, four cell phenotypes were predominant in patient-derived profiles, namely perforin^+^ T cell 8 (MAIT), CD38^+^ T cell 9 (CD8^+^ TEMRA-like), T cell 3 (γδ^+^ CD56^+^) and NK 3 (CD16^+^) cells. The frequencies of the respective host cell phenotypes were statistically higher in patient- compared to healthy control-derived *ex vivo* granulomas (**Fig. 3b**). The overall subset frequencies (irrespective of functional marker positivity) of T cell 9, but not that of the other three subsets, was significantly higher in patients than in healthy controls (**Fig. 3c**). Furthermore, and as depicted in a UMAP visualization, the cell subsets characterizing patients’ *ex vivo* granulomas partially shared the expression of perforin, CD38, as well as CD56 (**Fig. 3d**). Taken together, our data indicate that patient-derived granulomas are enriched in activated, cytotoxic NK and unconventional T lymphocytes.

**Figure 3.**
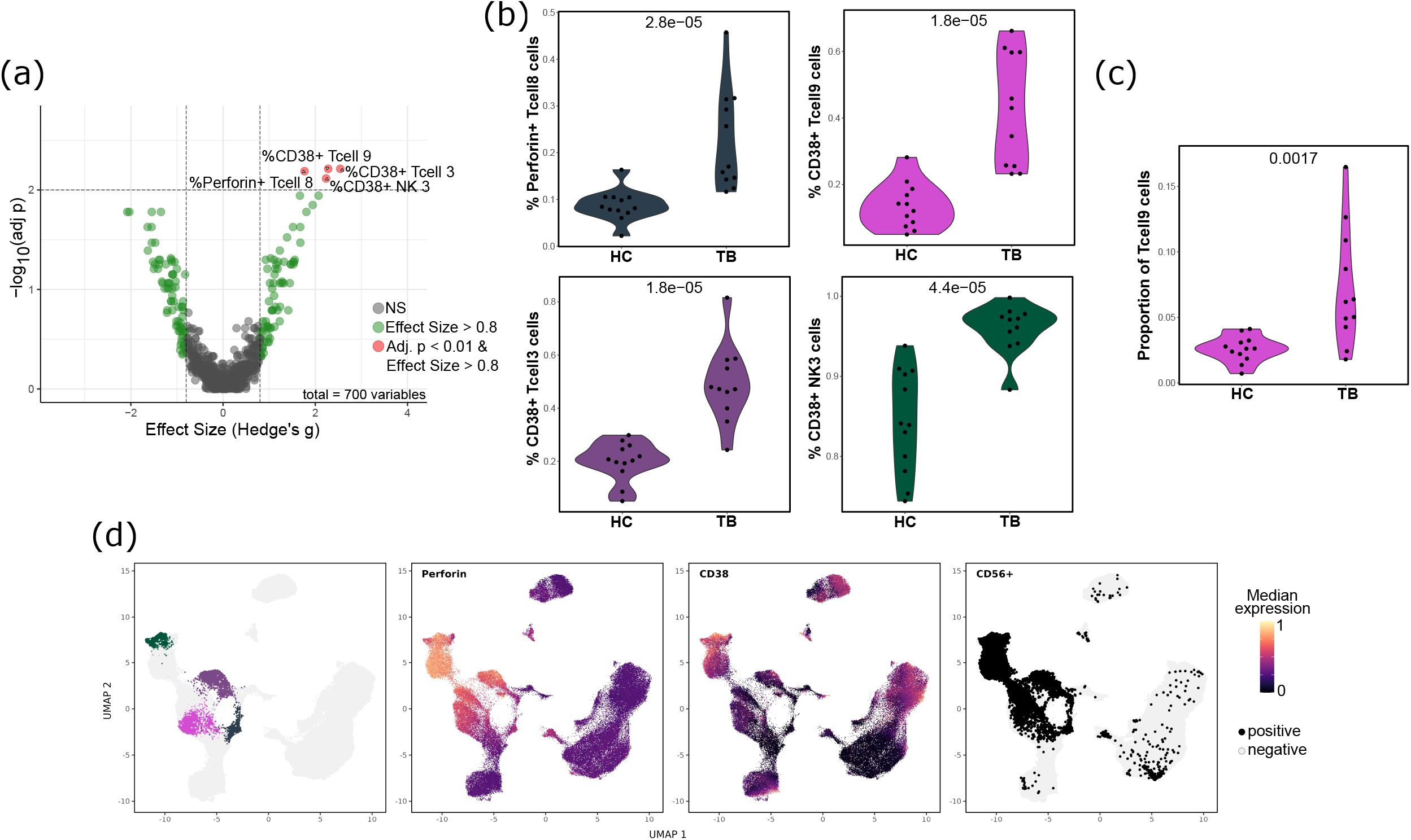
Cytotoxic, activated NK and (unconventional) T cells are significantly enriched in patient- derived granulomas. (a) Volcano plot (x: Hedge’s g effect size, y: false discovery rate (FDR) adjusted *p* values of pairwise Wilcoxon rank sum tests) of functional marker expression (patients vs healthy controls). (b) Frequency of host cell phenotype hits identified in (a) and, (c) frequency of T cell 9 subset (irrespective of functional marker positivity), split by disease group (inset *p* values from Wilcoxon tests). (d) UMAP visualizations of host cell phenotype hits identified in (a) (left) and, cells’ median expression of perforin (second left) and CD38 (second right) and CD56 positivity (right).

### Exposure to sub-mycobactericidal concentrations of granulysin drives *Mtb* to adopt a dormant-like state

We next investigated whether the host cell phenotypes enriched in patient-derived granulomas confer the increased prevalence of dormant-like *Mtb* following *ex vivo* infections. The frequencies of CD38^+^ NK 3 (CD16^+^) and T cell 3 (γδ^+^ CD56^+^) subsets showed a significant and strong positive correlation with the percentage of dormant-like *Mtb* (**Fig. 4a**). Given the shared expression of CD56 by these lymphocytic populations, we next depleted patients’ PBMCs of their CD56^+^ fraction and repeated the infections with *Mtb* to assess the role of CD56^+^ lymphocytes in the dormancy induction of *Mtb*. CD56^+^ cell depletion significantly reduced the percent dormant-like *Mtb* recovered from the *ex vivo* infections (**Fig. 4b**), while CD56^+^ cell complementation of CD56^+^-depleted specimens restored the dormancy level measured in untouched PBMCs (**Fig. S4A**). Flow cytometry analysis of the isolated CD56^+^ fraction revealed that it was mainly composed of NK cells (mean 77.9%), followed by NKT cells (17.3%) and a minor fraction of γδ T and MAIT cells (3.2% and 1.5%, respectively) (**Fig. 4c, Fig. S4B**).

**Figure 4.**
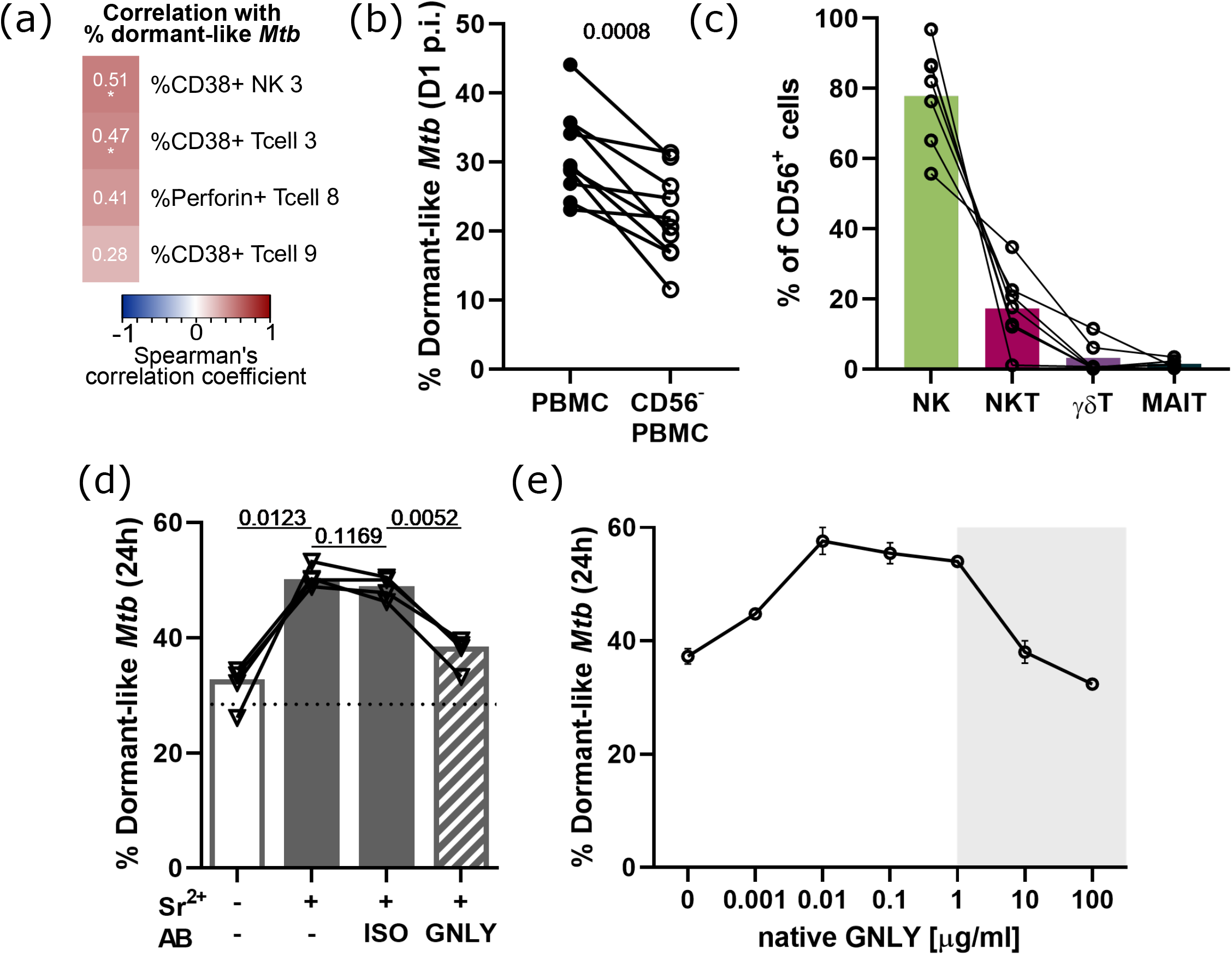
Exposure to sub-mycobactericidal concentrations of granulysin drives *Mtb* to adopt a dormant-like state. (a) Spearman’s correlation analysis between the frequency of dormant-like *Mtb* retrieved from *ex vivo* granulomas and the prevalence of indicated host cell phenotypes in the respective sample (asterisks for significant *p* values after Benjamini-Hochberg correction). (b) Percentage of dormant-like (Nile red^+^, auramine-O^-^) bacilli observed one day post-infection (p.i.) within *ex vivo* granulomas from TB patients depleted or not of CD56^+^ cells (n=10 independent patient specimens). Inset *p* value of a paired t test. (c) Composition of the CD56^+^ fraction depleted in b): NK (CD3^-^CD56^+^), NKT (CD3^+^), γδ T (CD3^+^TCRγδ^+^), MAIT (CD3^+^TCRVa7.2^+^); gating strategy provided as Fig.S4B. (d) Frequency of dormant-like *Mtb* upon 24h incubation with strontium-degranulated (+Sr^2+^) or not (-Sr^2+^) cell-free supernatants of CD56^+^ cells, in presence or absence of anti-granulysin (GNLY) or isotype control (ISO) antibodies (AB) (n=4 independent patient specimens). Dotted line indicates the *Mtb* dormancy level measured following incubation in cell-free strontium-containing medium. Inset *p* values from an ANOVA with Holm-Sidak’s multiple comparisons test. (e) Dose-response effect of native 9kDa granulysin^45^ on *Mtb* dormancy induction (based on auramine-O/Nile red staining). Error bars indicate the range and open circles the mean of two experimental replicates. Shaded area depicts the mycobactericidal concentrations based on Stenger *et al*. (1998)^31^.

Perforin, granzymes and granulysin are among the most abundant molecules contained in cytotoxic granules released by cytotoxic cells upon degranulation. In 1998, Stenger and colleagues ^31^ demonstrated the microbicidal activity of granulysin against a number of bacterial species, among which *Mtb*. This mycobactericidal activity of granulysin against extracellular and – in combination with perforin – intracellular *Mtb* has been confirmed independently^32, 33^. Moreover, granulysin has been shown to deliver granzyme B into bacteria, enabling its lethal effects^34, 35^. In that context, we next induced the degranulation of CD56^+^ cells with strontium and incubated *Mtb* with cell-free supernatants of strontium treated CD56^+^ cells. Degranulation-induced supernatants caused a significantly higher proportion of *Mtb* to adopt a dormant-like phenotype compared to the non-induced (control) supernatants. Addition of a granulysin-neutralizing antibody significantly reduced the induction of dormant-like *Mtb* (**Fig. 4d**). Moreover, incubation with native 9 kDa granulysin at sub- mycobactericidal concentrations resulted in the induction of dormant-like *Mtb* in a dose-dependent manner (**Fig. 4e**). Taken together, these data indicate that the delivery of granulysin by cytotoxic lymphocytes drives *Mtb* into a dormant-like phenotypic state.

## DISCUSSION

In this study, we unravel a role for granulysin in the induction of *Mtb* dormancy by combining an *ex vivo* granuloma model with TB endemic specimens and a high-dimensional mass cytometry approach. We describe PBMCs of healthy controls and TB patients to show comparable levels of reactivity to mycobacterial stimuli and not to differ statistically in their capacity to control *Mtb* growth *ex vivo*. Instead, patient-derived responses induce a dormant-like phenotype in *Mtb* within 24 hours of infection. We link this observation to the enrichment of patients’ specimens in cytotoxic and highly activated innate and innate-like CD56^+^ lymphocytes. Finally, we demonstrate that this dormancy induction in *Mtb* results from the exposure to sub-mycobactericidal concentrations of granulysin.

Our data indicate that the prevalence of *Mtb*-specific CD4 T cells does not translate into better control of *Mtb* growth *ex vivo*. This finding is supported by *in vivo* data of a clinical trial enrolling over 5,500 infants in South Africa, where the presence and responsiveness of mycobacteria-specific T cells was not protective against subsequent development of TB^36^. The estimation that 90% of *Mtb* infections do not result in clinical TB and the existence of so-called ‘resisters’ – heavily exposed individuals that never develop adaptive immune memory to *Mtb* – pinpoint innate immune mechanisms to be crucial in the early clearance of *Mtb*^37^. Interestingly, the CD56^+^ fraction in our depletion assays was mostly composed of innate and innate-like, unconventional T cells: NK, NKT, γδ T and MAIT cells. The presence of CD8^+^ lymphocytes – encompassing NK cells and CD8^+^ T cell subsets – has recently been found to confer protection in early non-human primate granulomas^21^. Activation of MAIT and γδ T cells has been observed in household contacts of patients with clinical TB, suggesting their involvement in the response to *Mtb* infection^38^. NK cells can directly sense microbial pathogens through Toll-like receptors^39^ and have been shown to readily react to mycobacterial stimulation^40^. Furthermore, NK cells are present in human TB lesions^40^ and their proportion increases in the peripheral blood during active TB compared to latently infected stages^18^, suggesting an involvement in TB pathogenesis. Here, the frequency of CD56^+^ and NK cells in particular did not differ statistically between patients and healthy controls (**Fig. S4c**). Nevertheless, our observations suggest that patients’ cytotoxic cells compared to those of healthy controls react more swiftly to release granulysin and drive this metabolic shift in *Mtb*. Further mechanistic studies are warranted to investigate if differential reactivity to direct mycobacterial sensing by innate (-like) cytotoxic cells exists between healthy controls and patients; or, whether additional cellular interactions or cytokines present in the granulomatous response are contributing to the enhanced activity of the patients’ specimens.

We previously linked the adaptation of a dormant-like phenotype by *Mtb* in our model to TNF- α^22^ and hypoxia^41^. Interestingly, hypoxia has been shown to induce higher levels of granulysin in human T cells^42^. Our results suggest that granulysin activity represents an additional immune-induced component within granulomas that drives *Mtb*’s stress response resorting to dormancy, only to resuscitate once the host gets immunocompromised.

Our approach has limitations, for PBMC-based models lack some cell types reported to constitute integral parts of *in vivo* granulomas, such as neutrophils and fibroblasts, whose presence might contribute to *Mtb* dormancy *in vivo* as well. Moreover, we cannot assert that immune cells present in *in vivo* granulomas harbor the exact characteristics of those circulating in the periphery. Nor do we claim that a one-week infection or a 24 hour assay mimic the pathophysiology of a chronic disease adequately. Nevertheless, we have previously demonstrated the clinical significance of *in vitro* granulomas in the context of *Mtb* dormancy and resuscitation^22^. The presented findings based on TB disease endemic specimens may serve to guide *in vivo* research and vaccine development in need of correlates of protection^43, 44^. In that context, it would be particularly interesting to investigate whether granulysin activity remains consistent across genetically distinct *Mtb* strains or lineages.

In conclusion, our data indicate that granulysin – delivered by innate (-like) cytotoxic cells upon mycobacterial infection – mediates a stress response that forces *Mtb* bacilli to enter a dormant-like phenotype. *Mtb* dormancy appears to be an important immune evasion mechanism counteracting the activity of this highly potent antimicrobial peptide. From a clinical perspective, tailored boosting of granulysin activity could prevent proliferation and favor *Mtb* clearance from the lung of infected individuals. In addition, cytotoxic functions and granulysin production in particular may be considered as relevant correlates of protection for TB vaccine development.

## METHODS

### PBMC isolation, cryopreservation & thawing

Peripheral blood mononuclear cell (PBMC) specimens were obtained from adult Tanzanian patients with active TB and healthy controls recruited within (i) a completed diagnostic study in Bagamoyo, Tanzania (TBChild project, IP.2009.32040.007) or (ii), patients only, an ongoing prospective cohort study in the Temeke District of Dar es Salaam, Tanzania (TBDAR). Both clinical study protocols were approved by the institutional review board of the Ifakara Health Institute (IHI), the Medical Research Coordinating Committee of the National Institute of Medical Research (NIMR) in Tanzania and the Ethikkommission Nordwest- und Zentralschweiz (EKNZ) in Switzerland. Some assays were set up using donors at Swiss blood banks in Bern or Basel (under informed consent).

PBMCs were isolated by Ficoll-Paque™ density-gradient centrifugation of fresh blood (clinical cohorts) or buffy coats (blood bank donors). PBMC rings were collected, washed twice and resuspended in RPMI-1640 medium with L-glutamine (RPMI), containing 10% DMSO and 40% fetal calf serum (heat-inactivated), before being aliquoted and cryopreserved in liquid nitrogen. For use, PBMCs were thawed, washed twice with RPMI supplemented with 10% FCS (complete medium) and 12.5 U/ml benzonase and rested over-night at 37°C, 5% CO_2_. The following day, PBMCs were counted by trypan blue exclusion method to confirm sample viability >90 %.

### *Mtb* H37Rv strain preparation

*Mtb* H37Rv was grown to a mid-exponential phase [OD_600_ ≈ 0.7] in Middlebrook 7H9 broth supplemented with 10% ADC (5% bovine albumin fraction V, 2% dextrose and 0.003% catalase), 0.5% glycerol and 0.05% Tween-80. Bacteria were washed with PBS + 0.1% Tween 80, pelleted at 3000×g (5 min) and resuspended in complete medium before dispersion by gentle water-bath sonication for 2 min (XUB5, Grant Instruments). Remaining clumps were pelleted at 260×g (5 min). The upper part of the supernatant was supplemented with 5 % glycerol (final), aliquoted and stored at -80°C. The concentration was determined through quantification of colony forming units (CFU) by 10-fold serial dilutions prepared in PBS + 0.1% Tween 80 and plated onto Middlebrook 7H11 agar plates supplemented with 0.5% glycerol and 10% OADC (0.05% oleic acid in ADC).

### PBMC stimulation

After overnight resting, PBMCs (2.5×10^5^ cells per condition in 200 μl final volume) were stimulated with *Mtb* whole cell lysate (10 μg/ml), *Mtb* peptide pool covering ESAT-6/CFP-10 sequences and parts of TB7.7 (5 μg/ml), *Mtb* soluble cell wall proteins (10 μg/ml), SEB (1 μg/ml, positive control) or left unstimulated. *Mtb* H37Rv whole cell lysate (NR-14822) and *Mtb* H37Rv soluble cell wall proteins (NR- 14840) were obtained through BEI Resources, NIAID, NIH. After 8 h of stimulation, one half of the supernatant was replaced with fresh complete medium containing brefeldin A (5 μg/ml). PBMCs were further incubated for 15 h before extra (ECS)- and intra (ICS)-cellular staining for flow cytometry. ECS was performed during 15 min at room temperature with mix of anti-CD27 (O323)-FITC (or anti-IgG1 (MOPC-21)-FITC isotype control) and CD14 (QA18A22)-APC/Cy7 antibodies (Biolegend). Cells were fixed with fixation buffer (Biolegend) and permeabilized with permeabilization wash buffer (Biolegend), following the manufacturer’s protocol. Subsequently, ICS was performed during 30 min incubation at room temperature with a cocktail of anti-CD3 (UCHT1)-PB, anti-CD4 (RPA-T4)-PE, anti- IFNγ (4S.B3)-PerCP and anti-CD8a (HIT8a)-APC antibodies (Biolegend).

### 3D *in vitro* granuloma model

3D *in vitro* granulomas were generated as previously described^22, 29^. In brief, overnight-rested PBMCs were resuspended in RPMI + 20% human serum (HuS; Pan Biotech, #P40-2701; heat-inactivated and filtered (Stericup, Millipore)) at 10^7^ cells per ml before infection, where appropriate, with H37Rv at an MOI of 1:400 (H37Rv:PBMC). 1.25×10^6^ infected/uninfected PBMCs were seeded across 48-well plates (Falcon) and embedded in an extracellular matrix (ECM) in a 1:1 (v/v) ratio before incubation at 37°C, 5% CO_2_ for 45 min. After incubation, the wells were topped up with an equal volume of RPMI + 20% HuS and further incubated until relevant time-points. The ECM was prepared by gentle mixing of 950 µl/ml of collagen (3mg/ml) (PureCol bovine type I collagen solution; Advanced BioMatrix), 50 µl/ml of PBS (10X), 4 µl/ml of fibronectin (1mg/ml) (from human plasma, 0.1% solution; Sigma-Aldrich) and 10 µl of NaOH (1N), followed by incubation at 4°C for up to 90 min. On day 1 and 4 post-infection, half of the culture supernatant was recovered and replace with fresh RPMI + 20% HuS. Granuloma formation was monitored by bright-field microscopy (Leica Thunder Imager Live Cell) on days 4 and 8 post-infection. When required, granulomas were treated with brefeldin A (5 μg/ml) 14 hours prior to intracellular cytokine assessment by CyTOF (see below). To recover host cells at relevant time point for downstream analyses (see below), supernatants were removed and the ECM was digested through the addition of 125 μl per well of collagenase (1 mg/ml). To enumerate CFU and assess *Mtb* dormancy, retrieved cell suspensions were lysed with Triton X-100 (0.1% final) for 25 min at room temperature.

### Depletion of CD56^+^ cells from PBMCs

The CD56+ fraction of PBMCs was depleted using Miltenyi’s human REAlease CD56 MicroBead kit (Miltenyi, 130-117-033), following manufacturer’s instructions. To assess depletion efficacy, individual fraction were stained with a mix of anti-CD4 (OKT4)-PB, CD8 (HIT8a)-BV510, CD56 (REA196)-FITC, CD19 (HIB19)-PE, CD14 (63D3)-PerCP, TCRγδ(B1)-PE/Cy7, TCRVa7.2 (3C10)-APC, CD3 (UCHT1)- APC/Cy7 antibodies for 15 min on ice. Anti-CD56-FITC was purchased from Miltenyi, all the other antibodies from BioLegend. PBMCs, CD56^+^-depleted PBMCs or CD56^+^-depleted PBMCs re-complemented with the depleted CD56^+^ fraction proportionally (1.25×10^6^ cells per well) were infected, embedded and retrieved as described above. *Mtb* dormancy was assessed 24 h post-infection, as described below.

### Sr2+ induced degranulation and incubation with *Mtb*

To induce degranulation, 10^5^ isolated CD56^+^ cells (see above) were incubated in complete medium containing 25mM strontium chloride (Sr^2+^) or equal volumes of 0.2 μm-filtered MiliQ water for 18 h (37°C, 5% CO_2_). Cells were pelleted and 100 μl of cell-free supernatants were inoculated with 2.62×10^4^ CFUs of *Mtb* H37Rv. Where indicated, supernatants were incubated with 0.2 μg/ml of an anti-granulysin antibody (Santa Cruz Biotechnology, sc-271119) or an isotype control (Biolegend, mouse IgG2a) for 1 h prior inoculation with bacteria. After 24 h of incubation, bacteria were processed for *Mtb* dormancy assessment, as detailed below.

### Effect of native granulysin on *Mtb*

Native granulysin (9 kDa) was extracted from YT Indy cells as previously detailed^45^ and tested for activity against *Listeria monocytogenes* (**Fig. S5**). 5×10^4^ CFU of *Mtb* H37Rv single-cell suspension in 7H9 were incubated with native granulysin at indicated concentrations for 24 h at 37°C. Dormancy was assessed as described below.

### Assessment of *Mtb* dormancy

On days 1 and 8 post-infection, bacteria in lysed cell suspension were pelleted at 6000×g (5 min) and inactivated with 1X CellFIX (BD) or 1X Fixation buffer (BioLegend) at room temperature (20 min), before storage at 4°C until staining. Inactivated bacilli were pelleted at 10000×g (5 min), resuspended in water, spotted on glass slides, air-dried and heat fixed at 70°C for at least 2 hours. Dual auramine-O/Nile red staining was performed as previously described^22^ (Arbués *et al*., 2020). Briefly, samples were stained with auramine-O (TB Fluorescent Stain Kit M, BD) for 20 min, decolorized for 30 s, stained with Nile red (10 μg/ml in ethanol, Sigma-Aldrich) for 15 min and counterstained with potassium permanganate (TB Fluorescent Stain Kit M, BD) for 2 min. Gentle washes with distilled water were performed between each step. After air drying, Vectashield mounting medium was applied and fluorescent bacteria (median number counted = 209, SD = 48) were manually enumerated by fluorescence microscopy (Leica DM5000 B).

### Quantification of *Mtb* load

On day 8 post-infection, 10-fold serial dilutions of lysed host cell suspensions were prepared in PBS + 0.1% Tween 80 in triplicate, plated on 7H11 agar, supplemented as described above, and incubated at 37°C, 5% CO_2_ for 4 weeks.

### CyTOF staining and acquisition

Retrieved host cells (see granuloma model) were stained using Standard Biotools’s products and following their protocol. All steps were performed at room temperature, unless otherwise indicated. In brief, cells were washed with Maxpar PBS (MP-PBS, Barium-free), stained in FcR block solution for 10 min and barcoded using CD45 antibodies for 15 min. Barcoded samples were washed twice with Maxpar Cell Staining Buffer (MP-CSB) before pooling and transferal to MDIPA tubes (max 3 million cells per tube), to which additional surface antibodies had been added (**Supplementary Table 2**). Cells were incubated with antibodies for 30 min, then washed twice before fixation with Maxpar fix I buffer for 25 min. Fixed samples were washed twice with Maxpar Perm-S Buffer (MP-PermSB), before intracellular staining for 30 min. Stained samples were washed twice with MP-CSB, fixed in 1.6% formaldehyde (Thermo Fisher Scientific; diluted in MP-PBS) for 10 min and stained with 125 nM Cell-ID intercalator-Ir (diluted in Maxpar Fix and Perm Buffer) for one hour. Samples were stored in 100 μl intercalator solution at -80°C until acquisition on a CyTOF XT. Prior acquisition, samples were washed with MP-CSB and CAS+ solution, strained and resuspended in CAS+ solution.

### CyTOF analysis

Samples of blood bank donors were included in all staining and acquisition runs and used for clustering. After cell identity assignment using FlowSOM (see below), the data set was reduced to include only samples of the Tanzanian clinical cohort (total of 2,288,786 cells).

#### Bead normalization, compensation and de-barcoding

Raw files were imported into R^46^ (version 4.2.0 (2022-04-22)) for bead normalization^47^, compensation (based on “CATALYST”‘s experimentally determined spillover matrix) and de-barcoding^48^ using “CATALYST”^49^. For more stringent de- barcoding, estimated separation cutoffs <0.2 were corrected to 0.2.

#### Manual gating

Debris, doublets and dead cells were gated out manually in FlowJo 10.6.1 (Java version 1.8.0_161-b12) (**Fig. S6a**). Gating was refined for each acquisition batch. Briefly, stable barium signal was selected, as previously described^50^. Subsequently, data was cleaned in consecutive gating of dense signal regions over time in beads (^140^Ce), residual, center, offset, width, event length, DNA1 (^191^Ir), DNA2 (^193^Ir), Live/Dead stain (adjusted from Standard Biotools).

#### Batch correction

Live singlets were imported into R (version 4.2.0), concatenated, arcsinh transformed (co-factor of 5) and batch corrected using “cyCombine”^51^. Each acquisition was considered an individual batch and the sample origin (i.e. clinical or blood bank) was used as a co- variant. The earth’s movers distance (EMD) reduction was 0.62 and median absolute deviation (MAD score) 0.02, pre- and post-correction UMAP visualizations showed minimal variation (**Fig. S6b**).

#### Transformation

Batch-corrected, arcsinh-transformed data was imported into R for 99.9^th^ percentile normalization. Markers CD45 and CD66b were removed.

#### FlowSOM clustering

Batch-corrected, arcsinh-transformed and normalized data was clustered using “FlowSOM”^52^. A hierarchical approach was applied (Level 1 and 2 clustering). At level 1, major cell types (i.e. T, B, NK, Myeloid cells) were separated based on the expression of lineage markers CD3, CD4, CD8a, CD19, CD20, HLADR, CD56NCAM, CD16, CD14, CD11c, CD33, CD 40. A 10×10 grid and 15 meta clusters were used. At level 2, further clustering was applied within each cell type based on cell-type specific lineage markers: T cells (CD3, CD4, CD8a, CD27, CD28, CD45RA, CD45RO, CD56, CD161, TCRgd, CCR4, CCR6, CXCR3, CXCR5; 10×10 grid and 12 meta clusters), B cells (CD11c, CD19, CD20, CD27; 8×8 grid and 4 meta clusters), NK cells (CD56, CD57, CD16; 8×8 grid and 3 meta clusters), myeloid cells (CD11c, CD14, CD16, CD38, HLA-DR; 8×8 grid and 6 meta clusters). Number of meta clusters was estimated using ‘MetaClustering’ (consensus) in the “FlowSOM” package and adjusted where necessary.

#### Marker positivity

Marker positivity cutoffs were estimated per sample using a silhouette-scanning approach in “MetaCyto”^53^. Overall median cutoffs were used as a basis for manual corrections.

#### UMAP visualizations

UMAPs were generated using the R package “umap”^54^ based on all the clustering markers used in “FlowSOM” and with the parameters n_neighbors = 15 and min_dist = 0.3.

#### Downstream analyses

Principal component analysis was performed on host cell phenotypes that were common across all samples (640 out of 700) with “factoextra”^55^. Fig. 2d was generated using the R package “Seurat”^56^. The volcano plot in Fig. 3a was generated based on all 700 host cell phenotypes using “EnhancedVolcano”^57^ and functions from Glass *et al*. (2023)^58^.

## ACKNOWLEDGMENTS

We would like to express our gratitude to all patients and controls who volunteered to take part in the studies. Additional acknowledgments to Dr. Sònia Borrell and the entire biosafety level 3 laboratory team at Swiss TPH for their support and to PD Dr. Amanda Ross for statistical advice. This research was funded by the Swiss National Science Foundation (grant numbers 197838 and 177163). Clinical samples used in this project were collected as part of the TBCHILD project (IP.2009.32040.007), funded by the European and Developing Countries Clinical Trials Partnership (EDCTP) and the TBDAR cohort. Calculations were performed at sciCORE (http://scicore.unibas.ch/) scientific computing center at University of Basel. Figure 1a was created with BioRender.com.

## AUTHOR CONTRIBUTIONS

Conceptualization, D.P., S.S.; Methodology, D.P., A.A., S.S., J.M.S.; Software, E.F.M., S.S.; Validation, A.A., S.S.; Formal Analysis & Visualization, S.S., E.F.M., D.P.; Investigation, S.S., D.P., A.A.; Resources, J.C., O.A.H., M.W., M.M., A.T., E.M., F.H., H.H., M.S., J.H.; Data curation, S.S., D.P.; Writing – Original draft, S.S., D.P.; Writing – Review & Editing, S.S., D.P., A.A., E.F.M, J.M.S., O.A.H., M.W., K.R., S.G., J.C., H.H., F.H.; Supervision, D.P., A.A.; Project administration, D.P.; Funding acquisition, D.P., K.R., S.G., J.F.

## COMPETING INTERESTS

J.M.S. and J.C. are full-time employees at Novartis.

All other authors have no competing interest to declare.

