## Supplementary Materials for "Granulysin antimicrobial activity promotes dormancy in *Mycobacterium tuberculosis*"

### SUPPLEMENTARY TABLES

Supplementary table 1: Characteristics of specimens used for the CyTOF immune profiling

| Group | % Female | Median Age (years) |
|---|---|---|
| **Healthy controls** (n=12) | 8.3% (1/12) | 21 |
| **Clinical TB** (n=11) | 23.3% (3/11) | 35 |

Supplementary table 2: CyTOF marker panel

| Target (clone) | Label | Used for | |
|---|---|---|---|
| | | FlowSOM clustering | Functional assessment |
| **CCR4** (L291H4) | 152Sm | YES | YES |
| **CCR6** (G034E3) | 141Pr | YES | YES |
| **CCR7** (G043H7) | 167Er | YES | YES |
| **CD11c** (Bu15) | 147Sm | YES | YES |
| **CD123** (6H6) | 143Nd | | YES |
| **CD127** (A019D5) | 176Yb | YES | YES |
| **CD14** (63D3) | 168Er | YES | |
| **CD16** (3G8) | 148Nd | YES | |
| **CD161** (HP-3G10) | 151Eu | YES | |
| **CD19** (HIB19) | 144Nd | YES | |
| **CD20** (2H7) | 171Yb | YES | |
| **CD25** (BC96) | 153Eu | YES | YES |
| **CD27** (O323) | 154Sm | YES | YES |
| **CD28** (CD28.2) | 160Gd | YES | YES |
| **CD294** (BM16) | 166Er | | YES |
| **CD3** (UCHT1) | 170Er | YES | |
| **CD33** (WM53) | 169Tm | YES | YES |
| **CD38** (HB-7) | 161Dy | YES | YES |
| **CD4** (RPA-T4) | 145Nd | YES | |
| **CD40** ( 5C3) | 142Nd | YES | YES |
| **CD45RA** (HI100) | 150Nd | YES | |
| **CD45RO** (UCHL1) | 149Sm | YES | |
| **CD56** (NCAM16.2) | 163Dy | YES | |
| **CD57** (HNK-1) | 155Gd | YES | |
| **CD69** ( FN50) | 113Cd | | YES |
| **CD80** ( 2D10.4) | 162Dy | | YES |
| **CD8a** (RPA-T8) | 146Nd | YES | |
| **CXCR3** (G025H7) | 156Gd | YES | YES |
| **CXCR5** (J252D4) | 158Gd | YES | YES |
| **GM-CSF** (BVD221C11) | 159Tb | | YES |
| **Granzyme B** (GB11) | 198Pt | | YES |
| **HLA-DR** (LN3) | 173Yb | YES | YES |
| **IFN-γ** (B27) | 116Cd | | YES |
| **IgD** (IA6-2) | 174Yb | YES | YES |
| **IL-10** (JES3-9D7) | 165Ho | | YES |
| **IL-1β** (CRM56) | 209Bi | | YES |
| **IL-6** (MQ2-13A5) | 106Cd | | YES |
| **PD-1** ( EH12.2H7) | 175Lu | | YES |
| **Perforin** (B-D48) | 196Pt | | YES |
| **TCRγδ** (B1) | 164Dy | YES | |
| **TNF-α** (Mab11) | 114Cd | | YES |
| CD66b (G10F5) | 172Yb | *excluded from analysis* | |
| CD45 (HI30) | 089Y | *excluded after data clean-up* | |
| DNA1 | 191Ir | *used for data clean-up / manual gating* | |
| DNA2 | 193Ir | | |

| | | |
|---|---|---|
| Live/dead | 103Rh | |
| CD45 (HI30) *barcoding* | 110Cd | *used for multiplexing* |
| | 194Pt | |
| | 111Cd | |
| | 195Pt | |

Supplementary table 3: Differential host cell phenotypes at $p < 0.05$ and effect size > 0.8

| **Host cell phenotype** | **Hedge's g effect size** | **Adjusted *p* value** |
|---|---|---|
| % CCR6+ Bcell1 cells | -2.0305 | 0.0166 |
| % CCR6+ Bcell3 cells | -2.0783 | 0.0166 |
| % CD28+ Myeloid2 cells | -1.6320 | 0.0407 |
| % CD38+ Myeloid3 cells | 1.2306 | 0.0407 |
| % PD1+ Myeloid3 cells | -1.6371 | 0.0236 |
| % CXCR5+ NK1 cells | -1.1316 | 0.0486 |
| % CD38+ NK2 cells | 1.6759 | 0.0339 |
| % CD38+ NK3 cells | 2.2346 | 0.0077 |
| % CD11c+ NK3 cells | 1.9434 | 0.0142 |
| % CD80+ Tcell1 cells | -1.5504 | 0.0236 |
| % CD28+ Tcell10 cells | -1.3476 | 0.0166 |
| % Perforin+ Tcell10 cells | 1.6664 | 0.0114 |
| % GranzymeB+ Tcell10 cells | 1.3871 | 0.0299 |
| % CD27+ Tcell11 cells | -1.4712 | 0.0339 |
| % CD38+ Tcell3 cells | 2.5414 | 0.0061 |
| % CD38+ Tcell4 cells | 1.8066 | 0.0166 |
| % CD25IL2Ra+ Tcell7 cells | -1.5461 | 0.0166 |
| % CD38+ Tcell7 cells | 1.5155 | 0.0236 |
| % CD38+ Tcell8 cells | 1.6738 | 0.0236 |
| % CD11c+ Tcell8 cells | 1.0329 | 0.0486 |
| % CD69+ Tcell8 cells | 2.0715 | 0.0114 |
| % CD127IL7Ra+ Tcell8 cells | -1.5468 | 0.0339 |
| % Perforin+ Tcell8 cells | 1.7668 | 0.0065 |
| % CD38+ Tcell9 cells | 2.2796 | 0.0061 |

SUPPLEMENTARY FIGURES

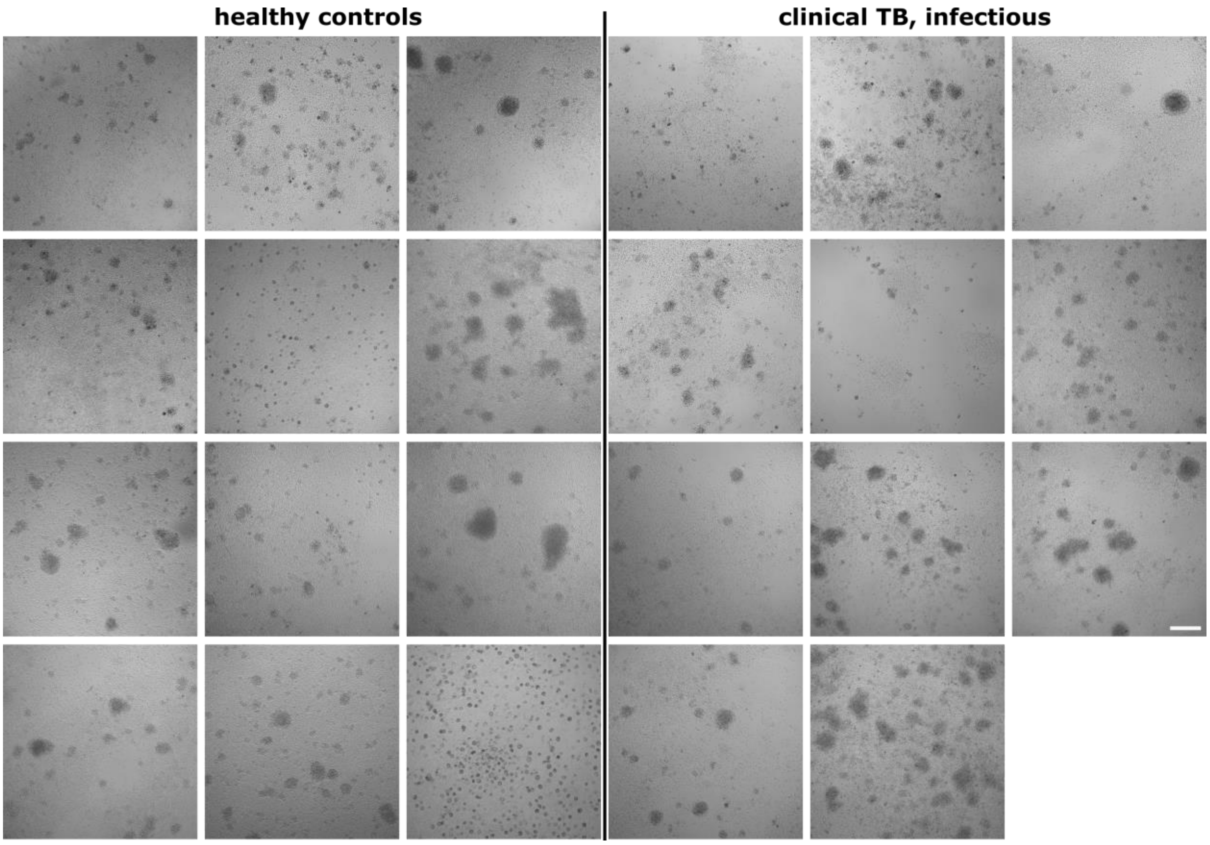


**Supplementary Figure 1. Monitoring of granuloma formation.** Bright field microscopy pictures of *ex vivo* granulomas formed eight days after infection and matrix embedding of PBMC samples obtained from each individual used in this study, split by disease status. Scale bar is 200 µm.

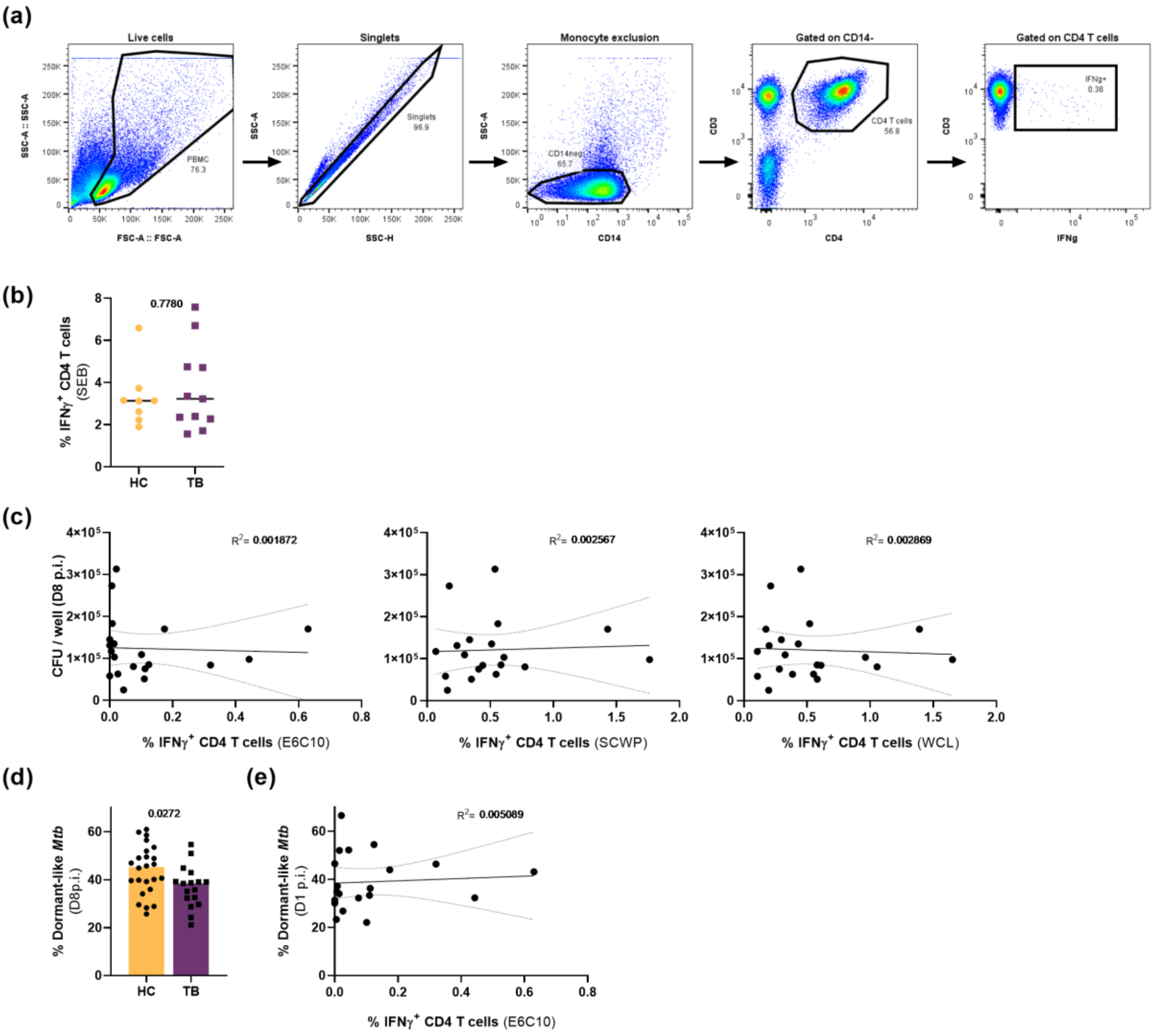


**Supplementary Figure 2. Assessment of *Mtb*-specific CD4 T cell response of PBMC specimens and correlation with bacterial read-outs.** (a) Gating strategy used to generate data presented in Figure 1B. (b) PBMC response to staphylococcal enterotoxin B (SEB) as positive control for sample integrity; lines indicate median. (c) Correlation between IFN-γ$^+$ CD4 T cells following overnight stimulation of PBMC samples and bacterial load recovered eight days p.i. from *ex vivo* granulomas formed with the respective PBMC samples. (d) Quantification of *Mtb* dormancy based on auramine-O/Nile red dual staining, eight days p.i.; bars depict the median. (e) Correlation between IFN-γ$^+$ CD4 T cells following overnight ESAT-6/CFP-10/TB7.7 peptide pool stimulation of PBMC samples and the frequency of dormant-like *Mtb* retrieved one day p.i. of the respective PBMC sample; all *p* values obtained from Mann-Whitney tests.

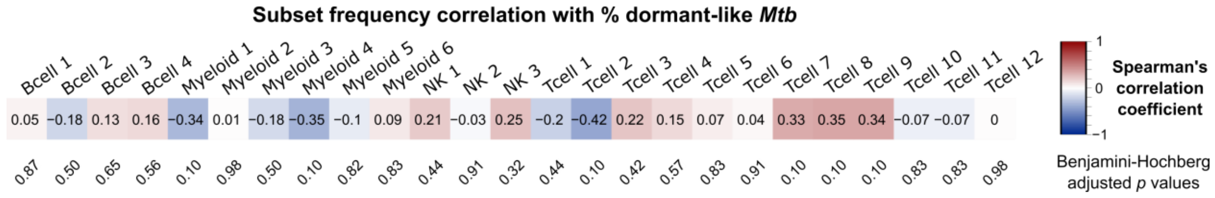


**Supplementary Figure 3. Correlation analysis of cell subset frequencies with *Mtb* dormancy**. Indicated subsets and frequency of dormant-like *Mtb* present in a given *ex vivo* granuloma sample. Numbers inside the squares indicate the Spearman's correlation coefficient. Numbers underneath in diagonal are Benjamini-Hochberg corrected *p* values.

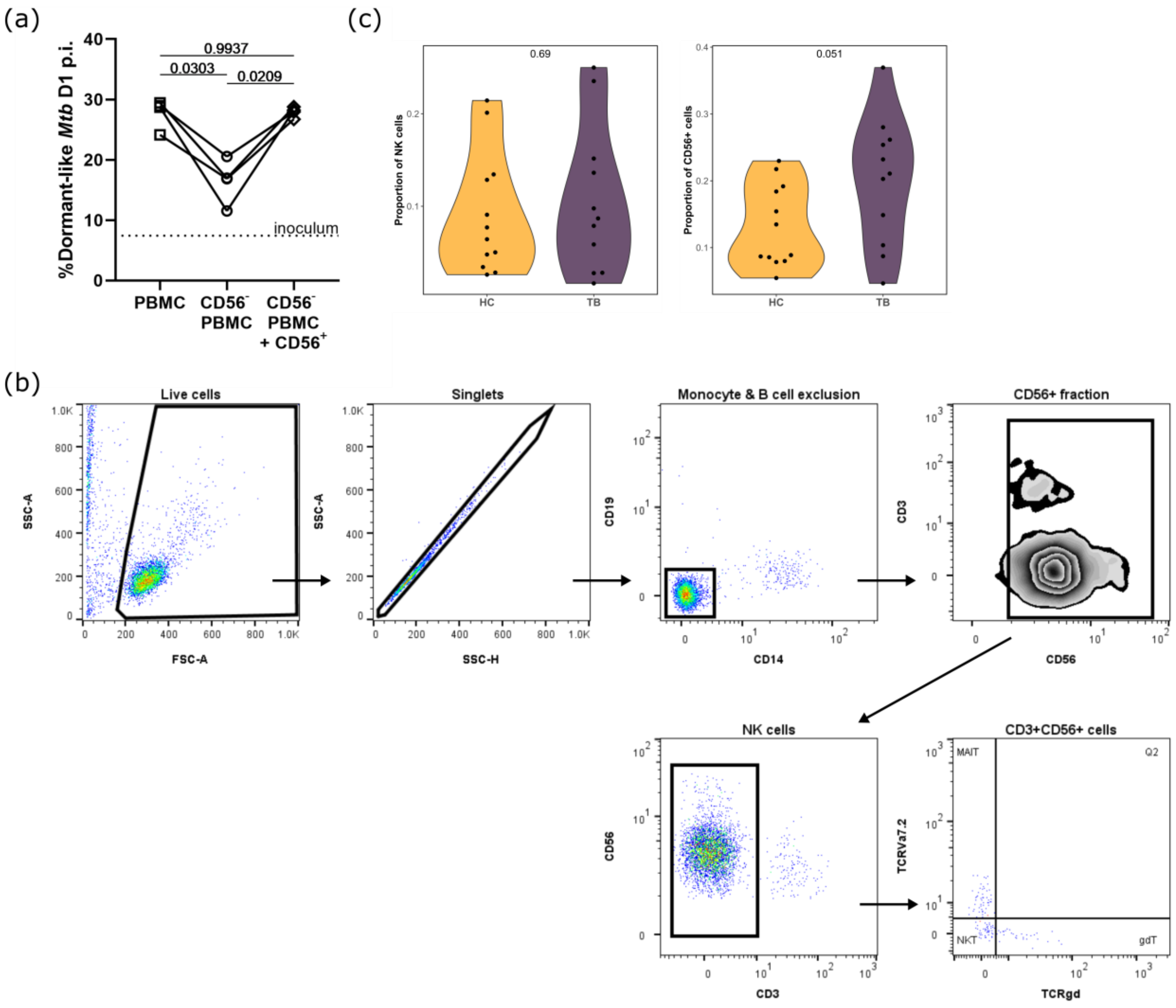


**Supplementary Figure 4. Depletion and reconstitution, gating strategy and frequency in PBMCs of the CD56$^+$ cell fraction.** (a) Percent dormant-like *Mtb* retrieved one day post-infection from *ex vivo* granulomas constituted of PBMCs,CD56$^+$-cell depleted PBMCs (CD56$^-$ PBMCs) or CD56$^-$ PBMCs that have been complemented with the removed CD56$^+$ fraction. Four independent patient specimens; *p* values indicate ANOVA tests with Turkey's multiple comparison correction. (b) Gating strategy applied to characterize the CD56$^+$ fraction used to generate the data presented in Fig. 4c. (c) Proportion of NK cells and CD56$^+$ cells split by group. Inset *p* values are from Wilcoxon tests.

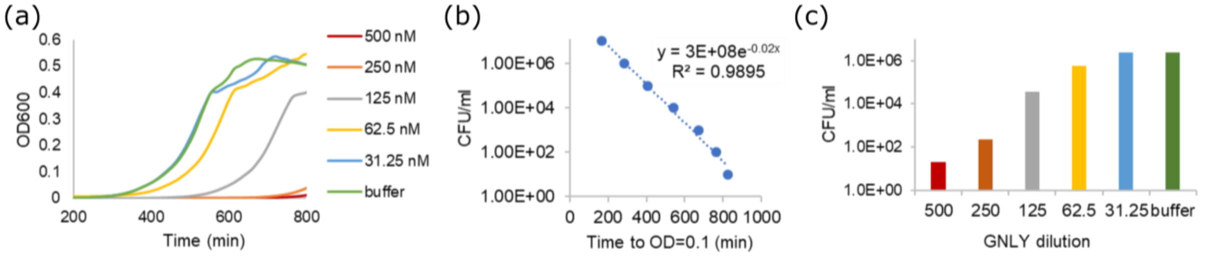


**Supplementary Figure S5. GNLY activity against gram+ bacteria.** *Listeria monocytogenes* 10403S were treated with indicated concentrations of native GNLY for 20 minutes at 37° C in assay buffer (50 mM NaCl, 20 mM Hepes, pH 7.4) before dilution (1:10) in brain heart infusion (BHI) broth. (a) Bacterial growth was monitored by overnight kinetics analysis at OD600 in a heat-controlled plate reader. CFU values were calculated using standard growth curves of serially diluted bacteria of known concentrations, plotted in (b), and are presented in (c).

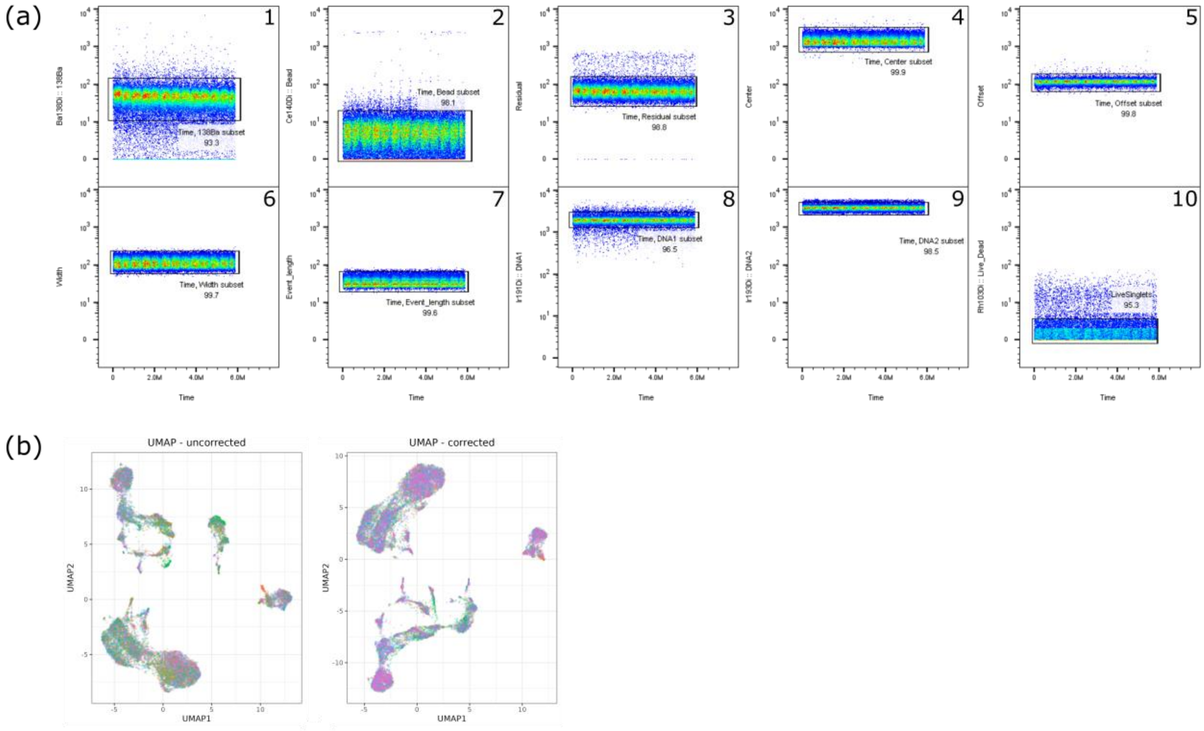


**Supplementary Figure 6. Live-cell gating and batch correction of the CyTOF data.** (a) Gating strategy applied for manual clean-up of CyTOF data. Numbers indicate consecutive steps. (b) Pre- and post-batch-correction UMAP visualizations. Colors indicate different staining and acquisition batches.
